# Preserving Vascular Integrity Protects Mice Against Multidrug-Resistant Gram-Negative Bacterial Infection

**DOI:** 10.1101/2020.02.18.955518

**Authors:** Teclegiorgis Gebremariam, Lina Zhang, Sondus Alkhazraji, Yiyou Gu, Eman G. Youssef, Zongzhong Tong, Erik Kish-Trier, Claudia V. de Araujo, Bianca Rich, Samuel W. French, Dean Y. Li, Alan L. Mueller, Shannon J. Odelberg, Weiquan Zhu, Ashraf S. Ibrahim

**Author notes:** Address Correspondence: Prof. Ashraf S. Ibrahim. David Geffen School of Medicine. Division of Infectious Diseases, Horbor-UCLA Medical Center. Currently employed by Merck (Dean Y. Li).

## Abstract

The rise in multidrug resistant (MDR) organisms portends a serious global threat to the healthcare system with nearly untreatable infectious diseases, including pneumonia and its often fatal sequelae, acute respiratory distress syndrome (ARDS) and sepsis. Gram-negative bacteria (GNB) including *Acinetobacter baumannii, Pseudomonas aeruginosa*, and carbapenemase-producing *Klebsiella pneumoniae* (CPKP), are among the World Health Organization and National Institutes of Health’s high priority MDR pathogens for targeted development of new therapies. Here we show that stabilizing the host’s vasculature by genetic deletion or pharmacological inhibition of the small GTPase ADP-ribosylation factor 6 (ARF6) increases survival rates of mice infected with *A. baumannii*, *P. aeruginosa*, CPKP pneumonia. We show that pharmacological inhibition of ARF6-GTP phenocopies endothelial-specific *Arf6* disruption in enhancing survival of mice with *A. baumannii* pneumonia, suggesting that inhibition is on target. Finally, we show that the mechanism of protection elicited by these small molecule inhibitors is by restoration of vascular integrity disrupted by GNB lipopolysaccharide (LPS) activation of TLR4/MyD88/ARNO/ARF6 pathway. By targeting the host’s vasculature with small molecule inhibitors of ARF6 activation, we circumvent microbial drug resistance and provide a potential alternative/adjunctive treatment for emerging and re-emerging pathogens.

## INTRODUCTION

Current global trends in healthcare-associated infections (HAIs) are characterized by the emergence and re-emergence of MDR pathogens (1). There are ~1.0 million HAIs in the US/year, which cause 70,000-100,000 deaths and $30B in costs (2). GNB are responsible for half of all HAIs. In particular, *Acinetobacter baumannii, Pseudomonas aeruginosa*, and carbapenemase-producing *Klebsiella pneumoniae* (CPKP) have emerged as predominant causes of MDR HAIs (3, 4). Some strains of these pathogens are resistant to all FDA-approved antibiotics and in many cases lead to serious complications (e.g., sepsis (5)), which are associated with high mortality and morbidity rates (6–8). Sepsis results from a dysregulated host immune response associated with production of a cytokine storm, which contributes to increased vascular permeability, tissue edema, organ failure, and death (9–12).

The past few decades provide evidence that development of resistance to antibiotics is inevitable. Thus, it is essential to develop novel treatments that do not rely solely on antibacterial action. Altering the patient’s own host response is one such alternative strategy for combating infectious diseases. Stabilizing the vasculature to prevent fluid leak, without compromising the beneficial innate immune system response, may allow the host ample time to clear the infection without causing organ failure. The activation state of ARF6, which is a convergence point in the signaling pathways of several inflammatory mediators and cytokines, plays a major role in controlling vascular permeability (13–16). When ARF6 is in its activated GTP-bound state, permeability is increased, and when it is in its inactive GDP-bound state, the vasculature is stabilized leading to decreased permeability. Proinflammatory agonists such as IL-1β, GNB lipopolysaccharides (LPS), and vascular endothelial growth factor (VEGF) activate ARF6, which in turn promotes the internalization of the adherens junction protein VE-cadherin and thereby increases endothelial permeability (13, 15, 16). We have shown that stabilizing the vasculature by reducing ARF6 activity is an effective strategy for reversing pathology and increasing survival rates in several preclinical models of inflammatory and vascular disease, including arthritis (15), LPS-induced endotoxemia (13), and diabetic retinopathy (16). Here, we show that stabilizing the vasculature through pharmacological inhibition of ARF6 activation by small molecules can increase survival rates and reduce pathology in preclinical models of MDR GNB infections.

## RESULTS AND DISCUSSION

### *Arf6* conditional knockout mice are resistant to *A. baumannii* pneumonia

We previously showed that mice infected with MDR *A. baumannii* die from endotoxemia caused by LPS activation of <u>T</u>oll Like <u>R</u>eceptor-<u>4</u> (TLR4) since TLR4-deficient mice are completely protected from *A. baumannii* bacteremia (17). We hypothesized that triggering of TLR4 by LPS during *A. baumannii* infection leads to activation of ARF6, resulting in increased virulence by compromising vascular integrity. To test this hypothesis, we compared the virulence of *A. baumannii* HUMC1 (a strain resistant to all antibiotics except colistin) (18, 19) in wild-type versus *Arf6* conditional knockout mice using pulmonary infection. *Arf6* conditional knockout mice were significantly more resistant to *A. baumannii* pneumonia with 40% overall survival after 21 days post infection versus 0% survival for mice carrying two active *Arf6* alleles (*Arf6*^*f*/+^) (Figure 1A). In another study, *Arf6*^*f*/+^ or *Arf6* conditional knockout mice (*Arf6^f/−^; +/Tie2-cre*) were infected with *A. baumannii* via inhalation and sacrificed for lung bacterial burden 4 days post infection. *Arf6* knockout mice had ~1.5 log_10_ lower *A. baumannii* burden than wild-type mice (Figure 1B). Collectively, these studies suggest that disruption of endothelial cell ARF6 allows the mouse immune system the opportunity to clear *A. baumannii* infection.

**Figure 1.**
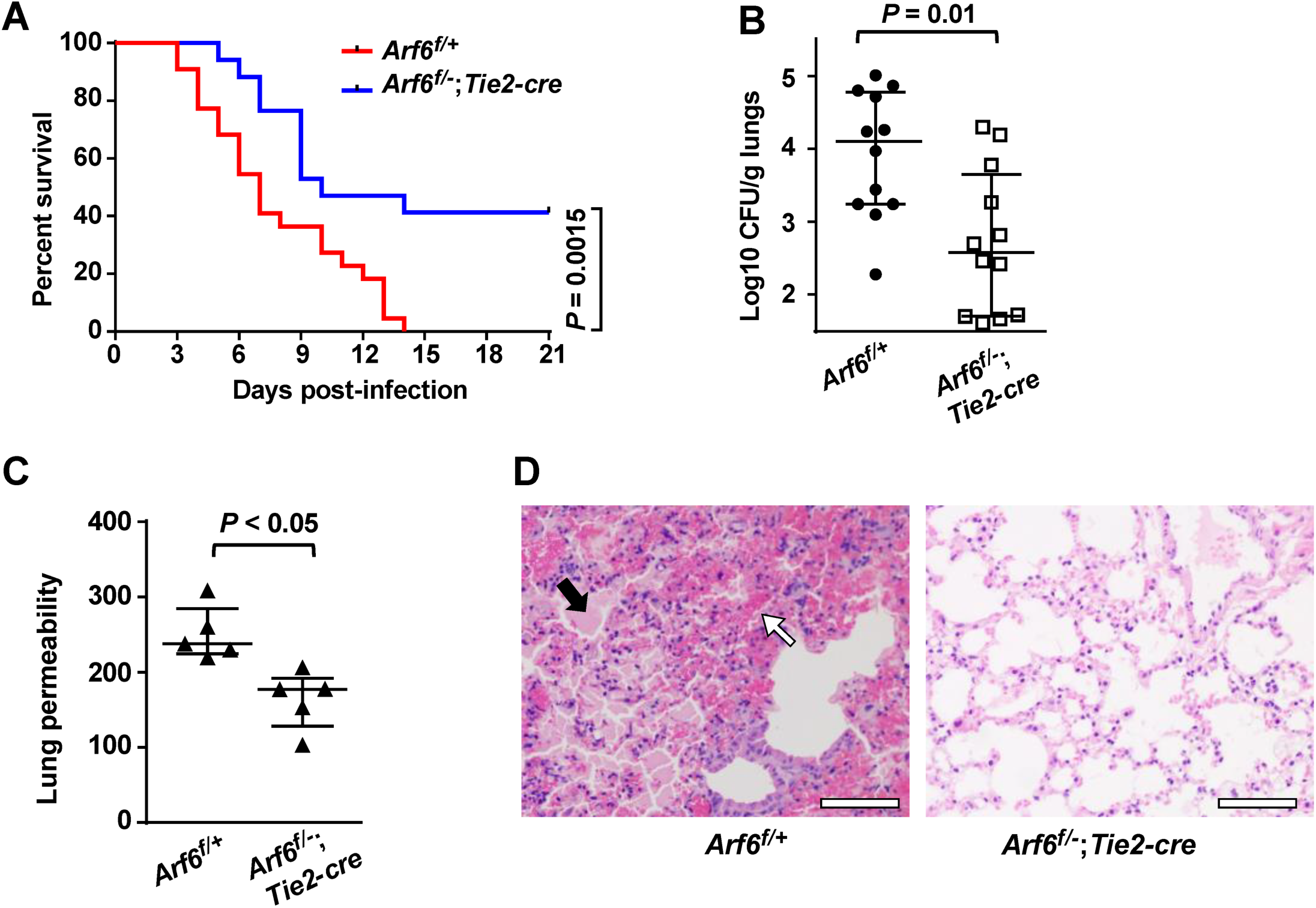
Endothelial cell *Arf6* deletion increases survival rates and reduces disease severity due to MDR *A. baumannii* pneumonia. (A) Survival of *Arf6*^*f*/+^ mice (n=22) or endothelial cell ARF6 null mice (*Arf6*^*f/−*^; *Tie2-cre*) (n=17) with *A. baumannii* pneumonia. (B) Lung bacterial burden; (C) lung permeability and (D) histopathological examination of lungs with H&E harvested from *Arf6*^*f*/+^ or endothelial cell ARF6 null mice 4 days post infection with *A. baumannii* via inhalation. Black and white arrows in (D) denote tissue edema and hemorrhage in wild-type mice, respectively. Bars are 20 μM. Data in (B) and (C) are presented as the median + interquartile ranges.

To test whether enhanced resistance of *Arf6* knockout mice to *A. baumannii* pneumonia is due to enhanced vascular integrity of the lungs, we infected mice with *A. baumannii* as above and 3 days later intravenously injected them with Evans blue dye to assess lung permeability. As expected, *Arf6*^*f*/+^ mice showed increased vascular permeability by ~25% when compared to *Arf6* conditional knockout mice (Figure 1C). Histopathological examination of the lungs corroborated the permeability results, with *Arf6*^*f*/+^ mouse lungs demonstrating enhanced tissue edema with extensive hemorrhage compared to *Arf6* conditional knockout mice (Figure 1D).

### ARF6 inhibitors are protective against murine bacterial pneumonia

Because ARF6 activation promotes virulence of *A. baumannii*, we hypothesized that pharmacologic ARF6 inhibition could be used to target GNB infections by stabilizing the vasculature. Thus, we tested whether pharmacologic inhibition of ARF6 could increase survival of mice with MDR GNB pneumonia. Small molecule ARF6 inhibitors, NAV-2729, A6-4424, and A6-5093 were evaluated for efficacy in treating murine pneumonia due to GNB. NAV-2729 (IC50 = 1.5 μM) has been shown to have therapeutic benefit in alleviating animal models of diabetic retinopathy (16). A6-5093 is a new highly soluble prodrug of A6-4424 designed to enhance *in vivo* administration. When A6-5093 is administered *in vivo*, its lysyl group is rapidly cleaved to produce A6-4424 (S1 Figure A), an active ARF6 inhibitor with IC50 = 1.9 μM (S1 Figure B-C).

Immunosuppressed mice were infected with aerosolized *A. baumannii* HUMC1 (a virulent MDR strain of *A. baumannii* that immunocompetent mice are resistant to) (18, 19). Mice were treated with: 1) 30 mg/kg of NAV-2729 or A6-4424 (both dissolved in DMA/PEG and the dose chosen based on pharmacokinetic (PK) data (S1 Figure D-G); 2) 43 mg/kg of A6-5093 (dissolved in saline and equivalent to 30 mg/kg of A6-4424); 3) DMA/PEG (vehicle); or 4) 2.5 mg/kg bid of colistin. Vehicle-treated mice had a median survival time of 8 days and almost complete mortality by Day 15 post infection. In contrast, NAV-2729 or A6-5093 enhanced median survival time to >21 days with an overall survival of >80% by Day 21 post infection (Figure 2A). Although mice treated with A6-4424 had a median survival time similar to vehicle-treated mice, 21-day survival was 50% versus 5% in vehicle-treated mice. Interestingly, the enhanced survival of mice seen with the ARF6 inhibitors far exceeded survival seen with the current first-line therapy of colistin (Figure 2A).

**Figure 2.**
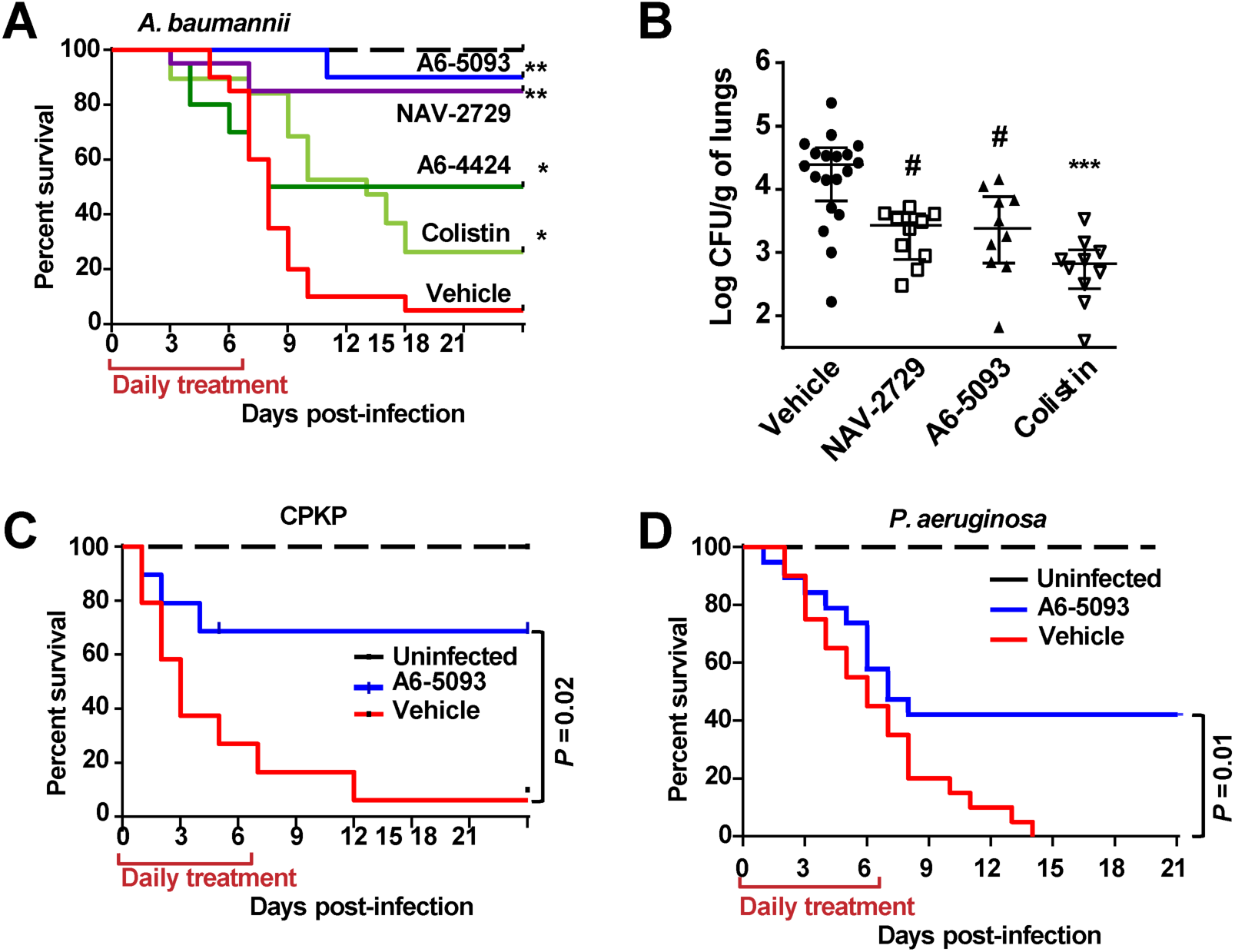
Pharmacologic inhibition of ARF6 increases survival rates and reduces disease severity due to MDR GNB infection. ARF6 inhibitors (A6-5093, NAV-2729, and A6-4424) increase survival (A, n=20 mice for all groups except A6-5093 and colistin which had 10 and 19, respectively) and reduce lung bacterial burden (B) of immunosuppressed mice (200 mg/kg IP cyclophosphamide and 250 mg/kg subcutaneous cortisone acetate given on Days −2 and +3 relative to infection) with MDR *A. baumannii* pneumonia. Lungs were harvested on Day +4, relative to infection. \**P*<0.05 vs. vehicle-treated mice and \*\**P*<0.02 vs. all others in (A) and #*P*<0.05 vs. vehicle-treated mice, \*\*\**P*<0.03 vs. all others in (B). ARF6 inhibitors increase survival rate of immunosuppressed mice infected with MDR CPKP (n=10/group) (C), or MDR *P. aeruginosa* (n=20/group) (D). Log Rank test for all analyses of survival and Wilcoxon Rank sum test for lung bacterial burden.

We investigated if the survival benefit correlated with reduction of bacterial burden in the lungs. Mice were infected and treated as above and sacrificed on Day 4 post infection. Mice treated with NAV-2729 or A6-5093 had at least 1-log_10_ reduction in lung bacterial burden compared to vehicle-treated mice (Figure 2B). Since *A. baumannii* HUMC1 is colistin sensitive (19), mice treated with colistin had a 2-log_10_ reduction of lungs bacterial burden versus vehicle-treated mice and 1-log_10_ reduction versus ARF6 inhibitors (Figure 2B). Thus, the decreased efficacy of colistin in survival studies is likely attributable to its toxicity (20).

We also evaluated the activity of A6-5093 against murine pneumonia caused by MDR *P. aeruginosa* or CPKP. Mice infected with CPKP or *P. aeruginosa* and treated with A6-5093 had 70% and 45% 21-day survival rates versus 10% and 0% survival rates for vehicle-treated mice, respectively (Figure 2C and 2D). Collectively, these results confirm the protective activity of ARF6 inhibitors against at least three MDR GNB, which are noted for causing HAI (21).

### ARF6 inhibitors enhance vascular integrity without affecting inflammation

We investigated the mechanism by which ARF6 inhibitors protect from GNB infection. First, we determined if any of our ARF6 inhibitors has a direct inhibitory or microbicidal effect on the three MDR bacteria under study. We performed minimal inhibitory concentration testing of the pathogen using either NAV-2729, A6-4424, or vehicle control. We observed no difference in *A. baumannii*, CPKP, or *P. aeruginosa* growth even at concentrations of 156 μM (S2 Figure), thereby indicating that the *in vivo* protection is likely due to an effect of the inhibitors on the host.

ARF6 activation leads to increased vascular permeability via internalization of the adherens junction protein VE-cadherin (13, 15, 16). Thus, we hypothesized that ARF6 inhibitors protect mice from GNB infection by maintaining vascular integrity. To investigate this possibility, we determined the effect of ARF6 inhibitors on *in vivo* organ permeability and the inflammatory response to *A. baumannii* pneumonia. Mice with *A. baumannii* pneumonia had an ~50% and 30% increase in lung and kidney permeability, respectively, when compared to uninfected mice. Treatment with either NAV-2729 or A6-5093 reduced the *A. baumannii-*mediated vascular permeability of lungs and kidneys to levels seen with uninfected mice (Figure 3A). These results were corroborated by histological examination of lungs harvested from mice infected with *A. baumannii*, which showed signs of pneumonia and tissue edema consistent with increased vascular permeability. Lungs harvested from mice infected and treated with either NAV-2729 or colistin showed more normal architecture with minimal to no signs of tissue edema (Figure 3B). Necrobiosis was evident in spleens taken from *A. baumannii-*infected mice without treatment or with those treated with colistin but not in spleens harvested from infected mice and treated with NAV-2729 (Figure 3B). The observed necrosis in spleens from colistin-treated mice is likely attributable to toxicity of colistin (20).

**Figure 3.**
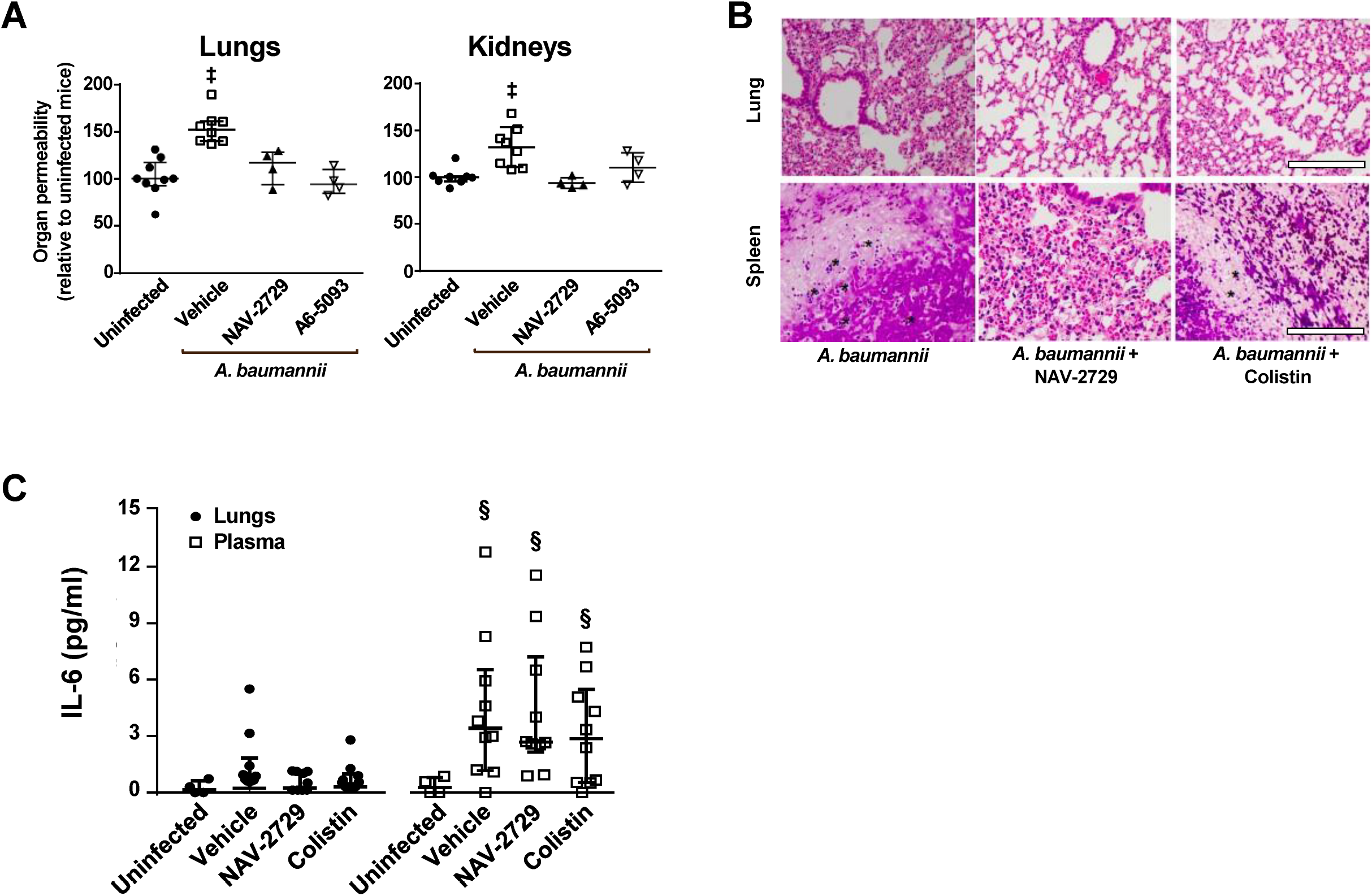
ARF6 inhibition reduces organ permeability and disease severity without affecting the inflammatory response to *A. baumannii* pneumonia. (A) ARF6 inhibitors (NAV-2729 and A6-5093) decreased permeability in the lungs and kidneys of *A. baumannii* HUMC1-infected mice. ^‡^*P*<0.05 vs. other treatments. (B) NAV-2729 (30 mg/kg) reduced severity of *A. baumannii* infection as shown by histopathological examination of lungs with H&E stain and spleen with Gram stain. * denotes necrobiosis in spleen harvested from vehicle-treated mice, and to a lesser extent colistin-treated mice, but not from mice treated with NAV-2729. Bars represent 100 μM. (C) IL-6 level in lungs and plasma of *A. baumannii* HUMC1-infected mice and treated with vehicle control, NAV-2729, or colistin. ^§^ *P*<0.02 vs. uninfected mice. Organs of neutropenic mice were harvested on day +3 for (A) and day +4 for (B) and (C), relative to infection. Data in (A) and (C) are presented as the median + interquartile ranges.

LPS-induced endotoxemia results in a robust immune response via TLR4-mediated NF-κB activation (17). To determine the effect of ARF6 inhibitors on the inflammatory immune response to *A. baumannii* pneumonia, we infected immunosuppressed mice and treated them as described above for 4 days. Mice were sacrificed and plasma and lungs were collected for determination of inflammatory cytokines. Due to the immunosuppression, which results in ~9 days of leukopenia (22), only IL-6 was detectable. No differences in IL-6 levels were observed in the lungs or plasma collected from vehicle-, NAV-2729-, or colistin-treated mice (Figure 3C). Collectively, these results show that ARF6 inhibitors likely protect mice from *A. baumannii* pneumonia by preserving vascular integrity without affecting the inflammatory immune response.

### Bacterial LPS induces vascular permeability via the MyD88/ARNO/ARF6-GTP pathway

Previously we showed that that TLR4 activation by LPS compromises vascular integrity via activation of the MyD88/ARNO/ARF6 pathway (13). It was also shown that lethality in mice infected with *A. baumannii* is mainly determined by the ability of the bacteria to activate TLR-4 through shed LPS. We hypothesized that GNB induces vascular permeability by activation of the MyD88/ARNO/ARF6 pathway through their LPS. To investigate this hypothesis, we studied if *A. baumanaii* and its LPS induce HUVEC vascular permeability *in vitro.* The virulent *A. buamanii* MDR HUMC1 and its cell-free supernatant (this strain sheds LPS (17)) induced HUVEC permeability equivalent to permeability seen with commercially available *E. coli* LPS, while the avirulent drug-sensitive strain ATCC 17978 cells and supernatant did not (this strain does not shed LPS (17)) (Figure 4A). To confirm that the enhanced HUVEC permeability observed following treatment with HUMC1 supernatant was due to LPS shedding, we removed LPS using polymyxin B agarose prior to incubating HUVECs with the supernatant (S3 Figure). Removal of LPS from the HUMC1 supernatant reduced permeability to uninfected HUVECs levels (Figure 4A), thus confirming the role of bacterial LPS in compromising vascular integrity.

**Figure 4.**
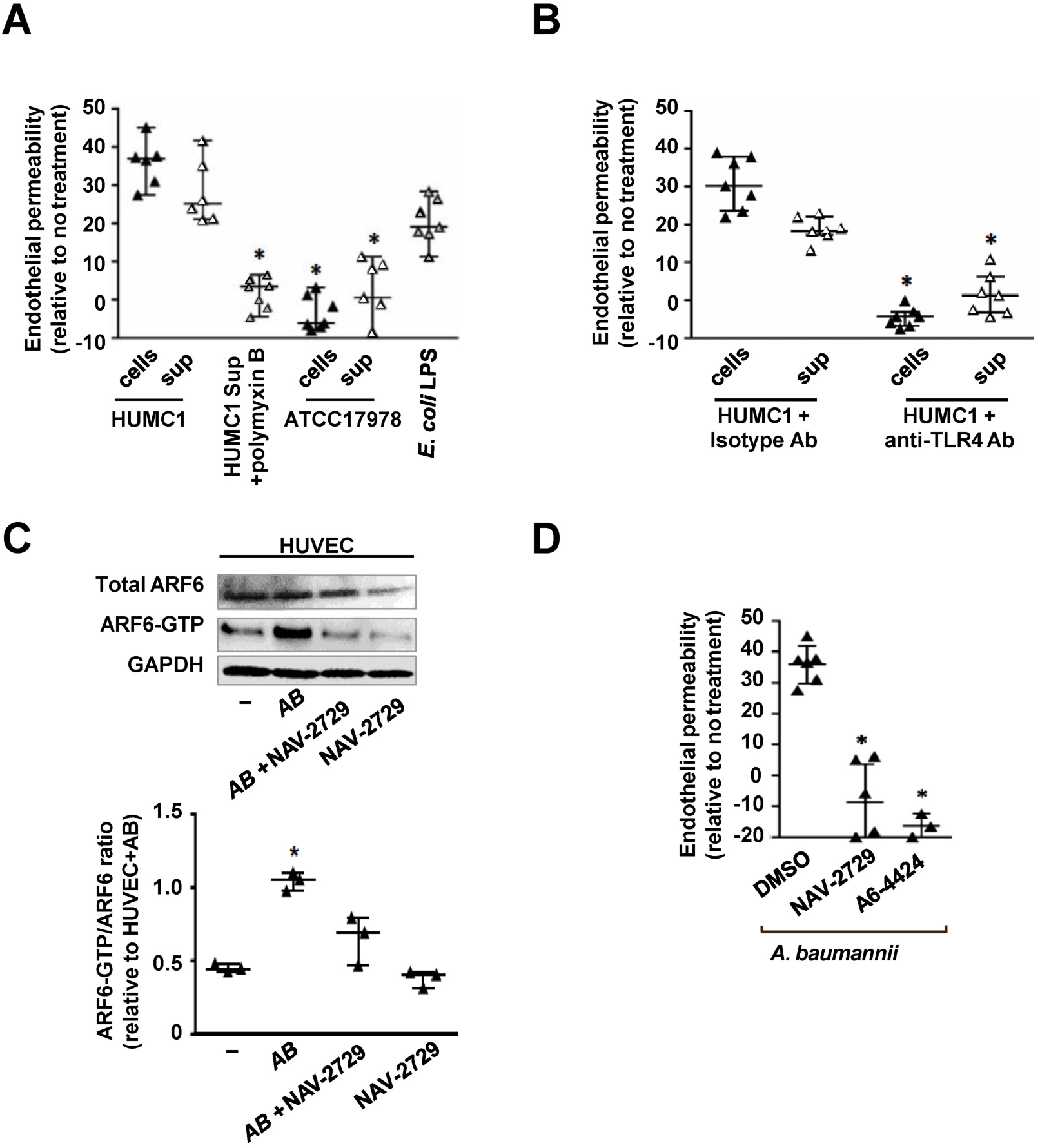
*A. baumannii*-mediated HUVEC permeability is induced by LPS-TLR4 signaling through ARF6 B. (A) *A. baumannii* HUMC1 (virulent MDR) cells and supernatants induce HUVEC permeability to a similar level as *E. coli* LPS. Removing LPS by polymyxin B blocks this induction. ATCC 17978 (avirulent, drug sensitive) cells and supernatants do not induce permeability. \**P* <0.005 vs. HUMC1, HUMC1 supernatant, or *E. coli* LPS. (B) A permeability assay was conducted with HUMC1 or its supernatant in the presence of 50 μg/ml of anti-TLR4 or isotype-matched control antibodies. \**P* <0.001 vs. isotype-matched antibody. (C), ARF6-GTP pulldown assays show that *A. baumannii* HUMC1 induces ARF6 activation and NAV-2729 blocks it. Quantification of the ARF6-GTP/total ARF6 ratio by densitometer (3 independent experiments) shows *A. baumannii* induces endothelial cell ARF6 activation by 2-fold, while NAV-2729 inhibits this activation. *\*P* <0.05 vs. all other comparators. (D) A HUVEC permeability assay was conducted with HUMC1 in the presence of 50 μM ARF6 inhibitors (NAV-2729 or A6-4424). \**P* <0.0015 vs. HUMC1 cells.

Mice with a TLR4 mutation or TLR4 knockout mice are resistant to *A. baumannii* infection (17). Thus, we hypothesized that blocking TLR4 would abrogate the ability of *A. baumannii* to induce HUVEC permeability. Indeed, anti-TLR4 antibodies completely blocked the ability of *A. baumannii* HUMC1 or its cell-free culture supernatant to induce HUVEC permeability (Figure 4B). Finally, by using ARF6-GTP pulldown assay (Figure 4C), we show that *A. baumannii* activated HUVEC ARF6 and NAV-2729 or A6-4424 reversed this activation and its sequelae of endothelial permeability (Figure 4D). Similarly, A6-5093 reduced CPKP- or *P. aeruginosa-*induced HUVEC permeability (S4 Figure).

ARF6 activation can be induced by multiple mechanisms including MyD88/ARNO (via TLR4 or the IL-1 receptor), the TNF receptor, or the VEGF receptor(11, 13–15). Since LPS-TLR4 interactions are critical for *A. baumannii* virulence, we hypothesized that activation of ARF6 occurs through stimulation of the MyD88/ARNO pathway. Consequently, downregulation of the expression of MyD88, ARNO, or ARF6 should protect HUVECs from *A. baumannii*-induced permeability. We used siRNA targeting MyD88, ARNO, or ARF6 to detect the contribution of these proteins to *A. baumannii-*induced vascular permeability. Downregulating the expression of MyD88 or ARNO by 70-85% (Figure 5A) resulted in ~50% reduction in HUVEC ARF6-GTP levels in HUVECs infected with *A. baumannii* (Figure 5B). As expected, transfecting HUVECs with siRNA targeting MyD88, ARNO, or ARF6 almost completely protected them from *A. baumannii*-induced permeability (Figure 5C).

**Figure 5.**
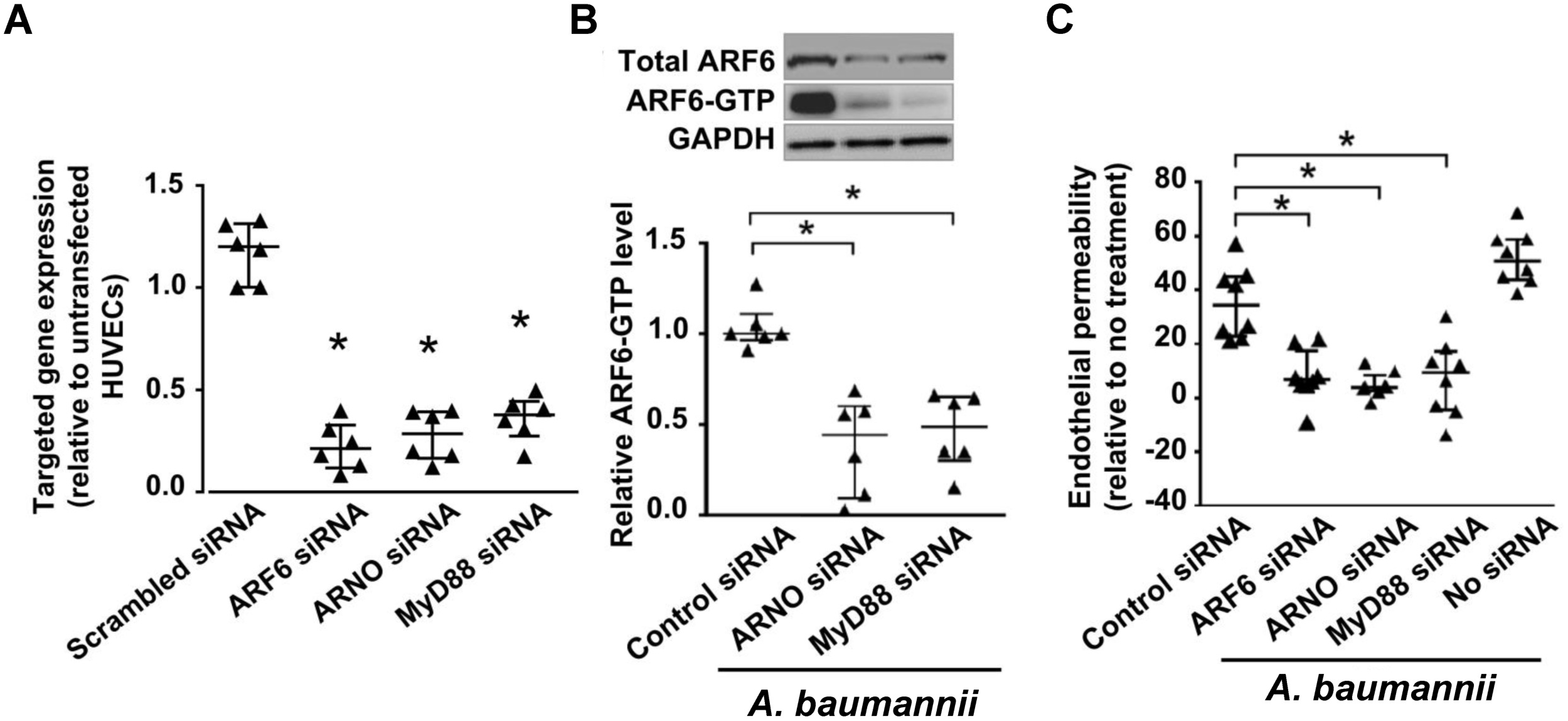
*A. baumannii*-mediated HUVEC permeability is induced through the MyD88/ARNO/ARF6 pathway. (A) Successful down regulation of gene expression (qRT-PCR) using siRNA constructs targeting ARF6, ARNO, or MyD88, in HUVECs. \**P* <0.001 vs. untransfected HUVECs (2 experiments). (B) (C) ARF6 expression as determined by qRT-PCR and Western blot (B) and *A. baumannii-*mediated permeability using FITC-dextran (C) of HUVECs transfected with siRNA constructs targeting MyD88, ARNO, ARF6 or scrambled sequence (control). \**P*<0.008 vs. control siRNA + *A. baumannii* (B), and *\*P* <0.002 vs. control siRNA + *A. baumannii* (C).

Collectively, these data show that vascular permeability is induced by GNB LPS via TLR4 activation of the MyD88/ARNO/ARF6 pathway and that ARF6 inhibitors minimize GNB-induced vascular leak *in vitro*.

### *A. baumannii* enhances HUVEC permeability by promoting internalization of VE-cadherin

We previously showed that LPS compromises vascular integrity by disruption of adherens junctions via internalization of VE-cadherin (13). To investigate whether a similar mechanism might be driving vascular permeability during *A. baumannii* infection, *A. baumannii* HUMC1-infected HUVECs were incubated with or without NAV-2729 and then immunostained with VE-cadherin antibody without permeabilization. *A. baumannii* reduced VE-cadherin surface expression at the cell-cell junctions (Figure 6A) and resulted in disrupting the monolayer by 25%. In contrast, addition of NAV-2729 to infected HUVECs preserved the surface expression of VE-cadherin and maintained the integrity of the monolayer to levels similar to those seen with uninfected HUVECs (Figure 6A-C). Thus, *A. baumannii* compromises vascular integrity by causing VE-cadherin internalization subsequent to ARF6 activation.

**Figure 6.**
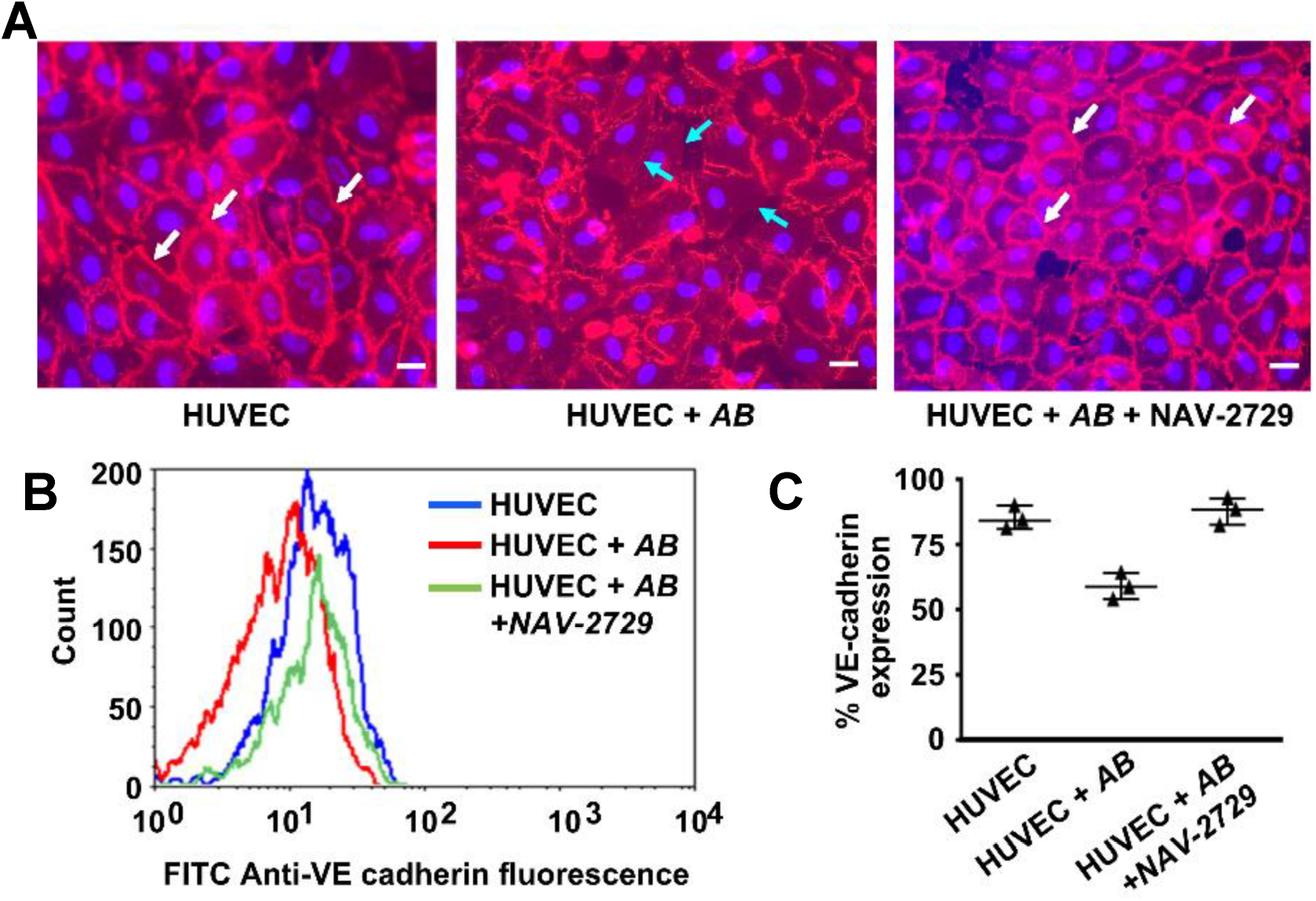
*A. baumannii* compromises HUVEC integrity by promoting internalization of VE-cadherin. (A) Confocal images of HUVECs incubated with *A. baumannii* HUMC1 (AB) with or without 50 μM NAV-2729 and stained with anti-VE cadherin antibody showing decrease in surface localization of VE-cadherin (red stain) at cell-cell junctions in the absence of NAV-2729 (cyan arrows) and restoration of junctional VE-cadherin with NAV-2729 to levels seen with uninfected HUVECs (white arrows). Cell nucleus is stained blue. Bars are 50 μM. (B) representative flow cytometry staining of surface expression VE cadherin and (C) its quantification from three independent experiments.

Based on these data and previously published results (13–16), we propose a model of MDR GNB virulence that relies on LPS binding to TLR4, which activates two independent pathways that diverge at MyD88: 1) a pathway that activates a robust immune response via NF-κB, which is characterized by an increase in inflammatory cytokines; and 2) a pathway that compromises vascular integrity via activation of ARF6 and internalization of VE-cadherin, resulting in excessive vascular leak, tissue edema, organ failure, and death. ARF6 inhibitors block vascular leak without affecting the LPS-induced inflammatory immune response, thereby allowing the immune system to fight the infection (Figure 7).

**Figure 7.**
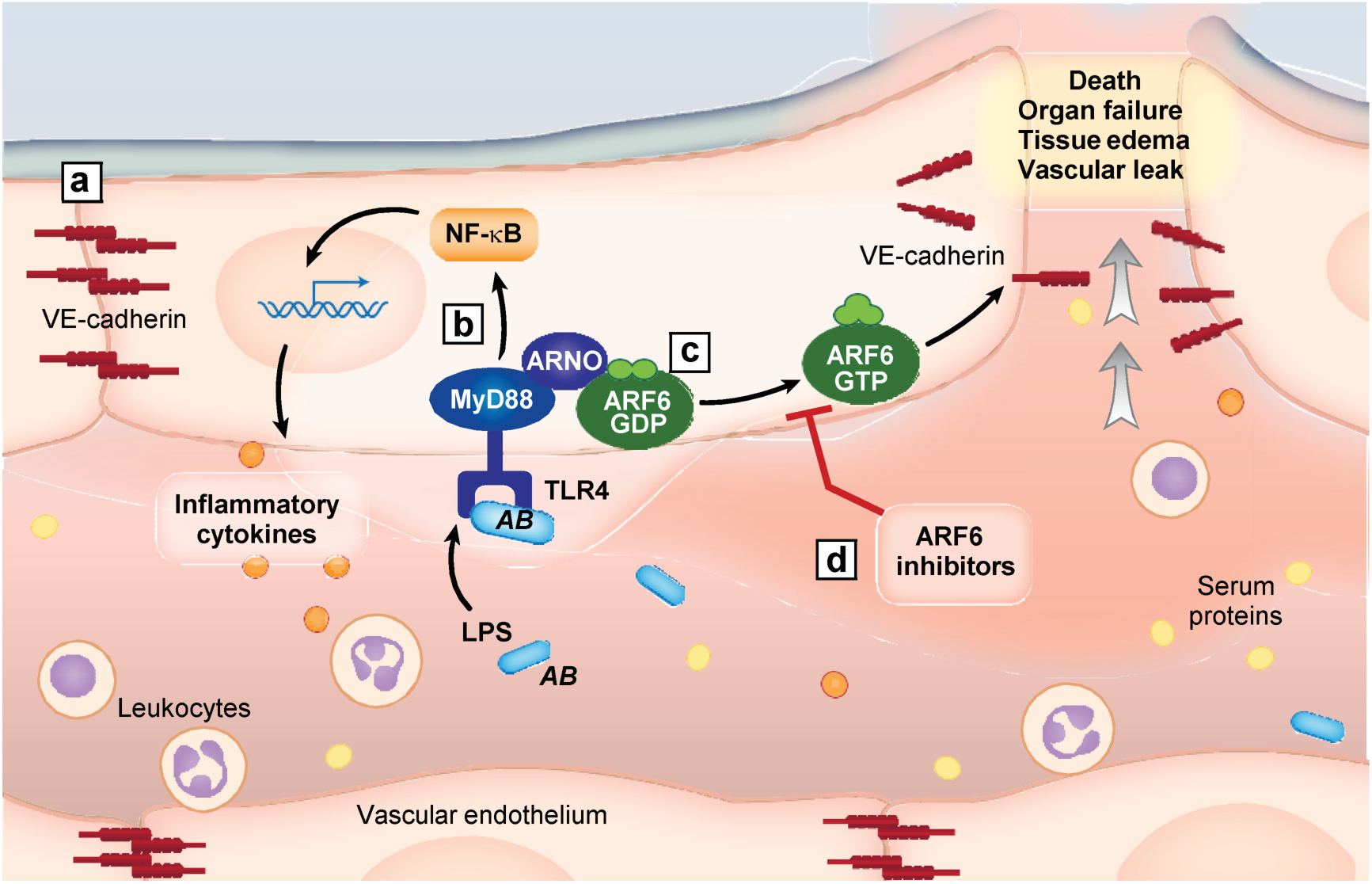
Model of GNB-induced vascular leak and the role of ARF6 inhibitor in preserving vascular integrity and increasing survival in bacterial infections. (A) In a normal host, the vasculature is held intact by VE-cadherin localized to the cell-cell junctions. (B) In bacterial sepsis (e.g. *A. baumannii*, AB), LPS-induced endotoxemia triggers a robust host inflammatory response by TLR4-mediated activation of MyD88/NF-κB required for clearing the infection. (C) Bacterial LPS also triggers ARF6 activation via MyD88/ARNO which leads to intercellular recruitment of VE-cadherin resulting in increased vascular leak, tissue edema, organ failure and ultimately death. (D) ARF6 inhibitors prevent ARF6 activation, which reduces VE-cadherin internalization resulting in the preservation of vascular integrity without affecting the inflammatory immune response.

By using ARF6 inhibitors to target the vascular leak that often accompanies severe infections, we have markedly increased survival rates in preclinical models of life-threatening infections of GNB. A primary advantage of this approach is that vascular leak is a host response to infection. Therefore, targeting it should not lead to antimicrobial resistance. This approach may prove to be applicable to any infection that activates MyD88/ARF6 (e.g., *Staphylococcus aureus* and fungal infections) (23, 24). Although our data show that monotherapy can have a marked effect on survival, in practice it is more likely that such an approach would be used as an adjunctive therapy to the limited drugs that are still in clinical use. Such combination therapies are the topic of active investigations.

## MATERIALS AND METHODS

### Organisms

*A. baumannii* HUMC1 is a MDR clinical strain with susceptibility to colistin (MIC = 2.0 μg/mL) (25). *A. baumannii* ATCC17978 is a carbapenem sensitive clinical strain (imipenem MIC = 0.25 μg/mL). Carbapenemase-producing *Klebsiella pneumoniae* (CPKP)-RM and *P. aeruginosa* are both MDR and obtained from patients at Harbor-UCLA Medical Center. Organisms were grown overnight in tryptic soy broth (TSB) at 37°C with shaking. Bacterial cells were passaged for 3 h in TSB, rinsed in phosphate buffered saline (PBS), and the final bacterial counts were determined using MacFarland standard at 600 nm. Susceptibility of the GNB to ARF6 inhibitors was assessed by the CLSI microdilution methods (26).

### Animal studies

#### Conditional *Arf6* knockout mice

C57BL/6 mice harboring the *Arf6*^*f*^ (*Arf6* with flanking LoxP sites (S5 Figure A) and *Arf6*^*−*^ alleles have previously been described (27). Endothelial-specific *Arf6*^*−/−*^ conditional knockout mice were generated by crossing *Arf6*^*f/f*^ mice with *Pdgfb-creERT2* mice. Excision of the *Arf6*^*f*^ allele in mice carrying the *Pdgfb-creERT2* transgene was accomplished by a single intraperitoneal injection of P1 mice with 40 μL of 1 mg/mL Tamoxifen in corn oil. *Arf6* excision in endothelial cells was verified by isolating lung endothelial cells as previously described (28) and performing a Western blot to assess ARF6 levels using anti-ARF6 antibody from Cell Signaling Technology (#3546) (S5 Figure B).

#### Infection with GNB

Male ICR mice (20-23 g) were obtained from Envigo. Mice (ICR or *Arf6* knockout mice) were immunosuppressed using 200 mg/kg cyclophosphamide (intraperitoneal [IP] administration) and 250 mg/kg cortisone acetate (subcutaneous administration) on Days −2 and +3 relative to infection. Mice were infected with aerosolized *A. baumannii* HUMC1 (18), or intratracheal instillation of CPKP (1.2 × 10^7^ cells) or *P. aeruginosa* (2.4 × 10^6^ cells). Treatment with ARF6 inhibitor (30 mg/kg once daily of active moiety given IP) started 3 h post infection. Infected mice treated with vehicle or those treated with 2.5 mg/kg of colistin given twice daily (bid) acted as controls. All treatments continued daily for seven days. The primary endpoints were time to moribundity. Secondary endpoints using organs harvested on Day +4 relative to infection, included tissue bacterial burden by quantitative culturing, inflammatory response, and histopathological examination using hematoxylin and eosin or Gram stain.

The inflammatory cytokine response (IL-1β, IL-6, and TNF-α) was determined using Cytometric Bead Array (BD Biosciences) per the manufacturer’s instructions.

Mouse organ permeability was determined by injecting Evans blue dye (EBD) (Sigma) 3 days after infection and harvested organs were processed as previously described (29, 30).

### Pharmacokinetic (PK) studies

#### Mouse

ARF6 inhibitor was administered to male C57BL/6 mice by IP injection. Mice were euthanized at various times after drug administration and blood was collected. Plasma was analyzed for presence of ARF6 inhibitor by a liquid chromatography/mass spectrometry (LC/MS) method. PK data were analyzed with Phoenix WinNonlin (Certara, Inc.)

#### Rat

ARF6 inhibitor was administered to male Sprague-Dawley rats by intravenous (IV) injection given over approximately 5 sec. Blood was harvested from indwelling arterial catheters at up to 10 time points over a 48-hr period after drug administration. Plasma was analyzed for presence of ARF6 inhibitor by an LC/MS method. PK data were analyzed with Phoenix WinNonlin (Certara, Inc.)

### Endothelial cells

Human umbilical vein endothelial cells (HUVECs) were collected by the method of Jaffe et al. (31) and passaged twice prior to using in assays. HUVEC propagation was at 37°C in 5% CO2 (32, 33). Reagents were tested for endotoxin using a chromogenic limulus amebocyte lysate assay (Charles River Laboratories), and the endotoxin concentrations were <0.01 IU/mL.

### Transwell permeability assay

HUVECs were seeded on 0.4 μm transwell inserts (Corning) coated with fibronectin (10-15 μg/mL in PBS) and fitted in 24-well plates (Costar). HUVECs were grown to confluency in M-199 medium without phenol red and then challenged with 10^7^ bacterial cells for 3 h at 37°C. Following incubation, 3 μL of 50 mg/mL FITC-dextran-10K (Sigma) was added to the top chamber and the amount of vascular leak was determined 1 h later by quantifying the concentration of the dye in the bottom chamber using florescence microplate reader at 488 nm.

To investigate the effect of stimuli or inhibitors on HUVECs permeability, the assay was repeated with or without 50 μM ARF6 inhibitors (A6 Pharmaceuticals), 50 μg/mL anti-TLR4 antibody or isotype control antibody (Biolegend), or 100 ng/mL *E. coli* LPS. To devoid bacterial supernatant of any shedded LPS, supernatants were treated with polymyxin B agarose beads.

### ARF6 pulldown assays

HUVECs were treated with *A. baumannii* HUMC1 with or without 50 μM ARF6 inhibitor for 4 h. The cells were washed with ice-cold PBS and lysed with ice-cold lysis buffer supplemented with protease and phosphatase inhibitor (Thermo Scientific, # 26186). Immunoprecipitation (IP) was performed with GST-GGA3-conjugated agarose beads (Cell Biolabs) (13). Protein level of GTP-bound ARF6 and total ARF6 in cell lysates was detected by Western blot with an anti-ARF6 antibody (1:1000 in 5% milk) (Sigma).

### HUVEC gene knockdown

HUVECs were trypsinized and resuspended in growth medium with 30 nM of siRNAs targeting MyD88 (SI00300909), ARNO (SI00061299), and ARF6 (SI02757286) using HiPerFect transfection reagent (Qiagen) (13) and incubated for 2 days at 37°C. The transfection process was repeated once more. All-Stars negative control siRNA (Qiagen) was used as a control. RNA from transfected HUVEC was isolated using RNeasy Plus Mini Kit (Qiagen) after incubation for 4 h with or without *A. baumannii* HUMC1. cDNA was synthesized using RETROscript Reverse Transcription kit (Thermo Fisher Scientific). qPCR reaction was prepared with Taqman Gene expression assay probes (Thermo Fisher Scientific) and TaqMan gene expression Master Mix (Thermo Fisher Scientific). The qPCR was carried out with a thermal-cycling program as follows: initial denaturing step for 10 min at 95°C, followed by 40 cycles of denaturing at 95°C for 15 s, and annealing/elongation at 60°C for 1 min. Human 18s *rRNA* gene probe (Hs00917508_m1) was used as a reference control. Real-time Probes of targeted genes were Human ARF6 (Hs01922781_g1), MyD88 (Hs00851874_g1), and ARNO (Hs00244669_m1). The comparative Ct method (ΔΔCT) was used for data analysis.

### VE-cadherin expression by confocal microscopy

HUVECs were seeded on coverslips that had been pre-coated with fibronectin in 24-well plates. HUVECs were incubated with *A. baumannii* HUMC1 for 4 h with or without 50 μM of ARF6 inhibitor, and then washed twice with PBS, fixed with 4% formaldehyde for 10 min. The cells were washed 4 times with PBS, and then blocked with 300 μl 50% human AB-serum (Sigma) for 10 min at room temperature. Mouse monoclonal VE-Cadherin antibody (BD Biosciences, # 555661) was added to the HUVECs (2.5 μg/mL) and held overnight at 4°C. Next, HUVECs were washed with PBS, counter stained with 2 mg/mL Alexa Fluor® 594 F(ab’)2 goat anti-mouse IgG (Thermo Fisher Scientific, 1:300 in 2% human serum-PBS) for 1 h at room temperature. The cells were stained with 1 μg/mL Hoeschst 33342 (ThermoFisher) in PBS for 10 min. HUVECs were washed twice with distilled water, and wet mounted on glass slides, before examining by confocal microscopy.

### VE-Cadherin quantification by flow cytometry

HUVECs were seeded in 6-well plate (Costar) pre-coated with fibronectin. HUVECs were incubated with *A. baumannii* HUMC1 for 4 h with or without 50 μM of ARF6 inhibitor, and then washed twice with PBS. HUVECs were detached from the plate with cellStripper dissociation reagent (ThermoFisher) that had been prewarmed at 37C for 10 min. M199 medium containing 20% FBS was added to each well and the cells collected. HUVECs were centrifuged, washed with PBS, and then fixed with 0.5 ml of 4% formaldehyde at room temperature for 15 min. The fixed cells were washed with PBS, blocked with 50% human AB-serum, and then incubated with 2.5 μg/ml mouse monoclonal VE-cadherin antibody (BD Biosciences, # 555661) overnight at 4°C. The next day, cells were washed with PBS and then incubated with Alexa Fluor® 488F(ab’)2 goat anti-mouse IgG secondary (ThermoFisher) (1:300 in 2% human serum-PBS from a stock of 2 mg/ml) for 1 h at 4°C. Cells were washed 3 times with 2% serum-PBS, and the cells were transferred to FACS tubes. Fluorescence of VE-cadherin was quantified by using a FACSCalibur (Becton Dickinson) instrument equipped with an argon laser emitting at 488 nm. Fluorescence data (10^4^ events) were collected with logarithmic amplifiers and % fluorescence was calculated using the CellQuest software.

### Ethics Statement

All procedures involving mice were approved by the IACUC of the Lundquist Institute for Biomedical Innovations at Harbor-UCLA Medical Center (Protocol number 30449), according to the NIH guidelines for animal housing and care. Moribund mice according to detailed and well-characterized criteria were euthanized by pentobarbital overdose, followed by cervical dislocation.

### Statistical analysis

Each *in vitro* experiment was carried out in triplicate and the experiment repeated at least twice. Survival was compared by the nonparametric log-rank. Categorical variables were compared with the Mann Whitney U test for unpaired comparisons. A *P* value < 0.05 was considered significant.

## Supplemental material

View Supplemental material

## Acknowledgments

This work was supported by the National Institutes of Health [R22-R33 AI119339 and R01 AI063503 to ASI; R01 HL077671 and R01 EY025342 to WZ; R01 HL130541, R01 AR064788, and R01 CA202778 to SJO; 1R43 HL127886-01, 2R44 HL127886-02A1, and 5R44 HL127886-03 to Navigen, Inc], and by UCLA CTSI grant UL1TR000124. The funders had no role in the study design, data collection, decision to publish or preparation of the manuscript.

We thank Ronell Lopez and Francisco Bautista for technical assistance with genotyping mice and helping with animal studies and Diana Lim for graphical assistance. We also thank Lise Sorensen and Yi Huang for technical assistance and Jacob Winter, Guy Zimmerman, and Michael Matthay for comments regarding the manuscript.

This work was presented in part at the ASM Microbe meeting held between June 16-20, 2016, Boston, MA and at the 2md International Symposium on Alternative to Antibiotics (ATA) held between December 12-15, 2016. Paris, France.

## Supporting information

**S1 Figure. A6-5093 is a prodrug small molecule inhibitor of ARF6.** (A) Structure of the prodrug A6-5093 and its active *in vivo* form (A6-4424) following cleavage of the lysyl group. (B) An ARF6 biochemical nucleotide exchange assay reveals an IC_50_ = 1.9 μM for the active form A6-4424. (C) A6-4424 inhibits ARF6 activation in a concentration dependent manner in a cellular ARF6-GTP pulldown assay. (D) Pharmacokinetic analysis of NAV-2729 (30 mg/kg IP) in mice (n=3/group). Dashed line represents biochemical IC_50_. (E) Changes in the plasma concentration of A6-4424 over time following intravenous injection of A6-5093 to rats. (F) Pharmacokinetics of A6-4424 after dosing of A6-5093 (30 mg/kg IP) in mice. (G) Tabulated PK properties of A6-4424 after IV administration of A6-5093 (top row) or A6-4424 (bottom row) to rats (3/group).

**S2 Figure. Effect of ARF6-GTP inhibitors on bacterial growth.** CLSI susceptibility testing showing lack of inhibition of *A. baumannii* (AB), CPKP, or *P. aeruginosa* (PA) by NAV-2729 or A6-4424.

**S3 Figure. Removal of shed LPS from MDR *A. baumannii* and inhibition of *A. baumannii*-induced ARF6 activation by small molecule inhibitor**. LPS concentrations in cell-free culture supernatant of MDR *A. baumannii* HUMC1 before or after mixing with polymyxin B agarose beads.

**S4 Figure. A6-5093 reduces CPKP- or *P. aeruginosa*-induced HUVEC vascular permeability.** Permeability was conducted using FITC-dextran. Data are from two independent experiments and presented as median ± interquartile ranges. Statistical analysis is by Mann-Whitney.

**S5 Figure. Strategy for endothelial cell deletion in C57BL/6 mice.** (A) Schematic of our approach to generate endothelial-specific knockout mice. P, loxP. Heavy lines represents the *Arf6* gene. Black rectangle within the gene represents the coding region. F, remaining recombined FRT sequence following the removal of the neo^r^ gene by flp recombinase. Adapted from Zhu *et al.* (27) (B) Western blot showing ARF6 expression in mouse lung endothelial cells for each of the four sibling genotypes.

